# Systematic prediction of genetic variance in a line development program

**DOI:** 10.64898/2026.11.16.767096

**Authors:** Rex Bernardo

## Abstract

To date, predicting the genetic variance (*V*_G_) in biparental crosses has been unreliable. My objective was to determine if predicting *V*_G_ via tabular analysis, which was developed assuming free recombination and having known *V*_G_ in founder crosses, is effective even if these unrealistic assumptions are unmet. With line X being developed from the P1/P2 cross, the *V*_G_ in X/Y was predicted as *V*_X/Y_ = λ_1_*V*_P1/Y_ + λ_2_*V*_P2/Y_, where λ_1_ was the parental contribution of P1 to X and λ_2_ was the parental contribution of P2 to X. Simulations involved four founder lines with biallelic marker loci and multiple alleles at quantitative trait loci. The *V*_G_ in each founder cross was estimated from simulated phenotypic values of *N* lines under varying heritability (*h*^2^). Non-founder lines were developed successively from the founders. Among the 774 non-founder crosses, the mean correlation between the true *V*_G_ and predicted *V*_G_ ranged from 0.49 (*N* = 50 and *h*^2^ = 0.20) to 0.70 (*N* = 400 to 1000 and *h*^2^ ≥ 0.50). The mean regression of predicted *V*_G_ on true *V*_G_ was 0.51–0.52. Having *N* = 100–200 lines was sufficient for estimating *V*_G_. Free recombination increased the regression of predicted *V*_G_ on true *V*_G_ to 0.87 and the correlation between the two to 0.64–0.93. Overall, the results suggested that tabular analysis is currently the most effective method for predicting *V*_G_. Further research is needed on combining *V*_G_ estimates in both founder and non-founder crosses when conducting tabular analysis in a breeding program.

**Plain Language Summary:** In plant breeding, the best sources of new varieties are populations that are both high performing and that have wide genetic variability. Predicting the mean performance of a breeding population is routine but predicting its genetic variance has been difficult. This simulation study found that the genetic variance in newer populations can be predicted systematically from the genetic variance in older populations, even when unrealistic but convenient genetic assumptions are relaxed. Routine application of tabular analysis to predict genetic variance will require methods to combine estimates of genetic variance in older breeding populations and in more-recent breeding populations.

**Core Ideas:**

- So far, plant breeders are unable to reliably predict the genetic variance (*V*_G_) in a cross between two lines.
- Tabular analysis can be used to predict *V*_G_ in new crosses from prior estimates of *V*_G_ in older crosses.
- Simulations showed that *V*_G_ predicted via tabular analysis was accurate, even when model assumptions are violated.
- We will need to see if tabular analysis will work well if some *V*_G_ estimates are unavailable in older crosses.

## 1 INTRODUCTION

Homozygous lines are routinely developed from biparental crosses in self-pollinated species, such as soybean [*Glycine max* (L.) Merr.], and in species for which hybrid cultivars are grown, such as maize (*Zea mays* L.). The biparental populations most useful in breeding combine a high mean performance (assuming higher values of the trait are favorable) and a large genetic variance (Bernardo, 2020; Melchinger et al., 1988).

Predicting the mean of a biparental cross is straightforward. If the mean of parent A is µ_A_ and the mean of parent B is µ_B_, then the mean of random homozygous lines derived from the A/B biparental cross is ½(µ_A_ + µ_B_) (Bernardo, 2020). The mean of random homozygous lines derived from the [A(A/B)] backcross population is ¾µ_A_ + ¼µ_B_. When the goal is to maximize testcross performance in a hybrid crop, the testcross mean of random recombinant inbreds is ½(µ_A/T_ + µ_B/T_), where µ_A/T_ and µ_B/T_ are the means of each parent when crossed to a common tester denoted by T (Melchinger, 1987). Both µ_A_ and µ_B_, as well as their corresponding testcross means, can be routinely estimated from field trials, with a mixed-model framework being useful for handling unbalanced data from trials conducted across different locations and years (Bernardo, 2020).

To date, however, the genetic variance (*V*_G_) in biparental crosses has not been reliably predicted. In this article, *V*_G_ refers to the additive genetic variance or the testcross variance expressed among homozygous lines. Early attempts to predict *V*_G_ relied on the pedigree distance or molecular marker distance between the parents of a biparental cross, but the resulting predictions were inaccurate (Cowen & Frey, 1987; Helms et al., 1997; Kisha et al., 1997; Manjarrez-Sandoval et al., 1997; Moser & Lee, 1994; Souza & Sorrells, 1991). In hindsight, the inability to predict *V*_G_ from pedigree or molecular marker distances is unsurprising: if such metrics were to be successful, then breeders should expect that the biparental crosses with the highest and lowest values of *V*_G_ should be the same regardless of the trait, and this expected result is contrary to breeders’ observations.

More-recent attempts to predict *V*_G_ have relied on genomewide marker effects estimated from empirical data (Bernardo, 2014; Mohammadi et al., 2015). This approach for predicting *V*_G_ involves simulating homozygous lines from a biparental cross, calculating the genotypic value of each simulated line from genomewide marker effects (Meuwissen et al., 2001), and calculating *V*_G_ as the variance among the genotypic values of the simulated individuals (Adeyemo & Bernardo, 2019; Lian et al., 2015; Neyhart et al., 2019; Wartha and Lorenz, 2024). Results from this approach have been mixed, with the correlations between predicted and observed *V*_G_ sometimes being low (Adeyemo & Bernardo, 2014; Lian et al., 2015) and sometimes being moderately higher (Neyhart et al., 2019; Wartha and Lorenz, 2024). Regardless of the correlation between the predicted and observed *V*_G_, the predicted *V*_G_ had a downward bias, which is expected because of shrinkage of genomewide marker effects toward zero (Meuwissen et al., 2001). The bias in the predicted *V*_G_ was as severe as –96% (Neyhart and Smith, 2019), and this downward bias is unacceptable because it penalizes the contribution of *V*_G_ in the predicted mean of the best progeny as expressed in the usefulness criterion (Melchinger et al., 1988).

An alternative approach is to predict *V*_G_ in new biparental crosses from estimates of *V*_G_ in prior biparental crosses. For example, the *V*_G_ in the A/B cross can be predicted as the mean *V*_G_ in all prior crosses that had A as one of the parents (e.g., A/C, A/D, A/E, etc.) and that had B as one of the parents (e.g., B/E, B/F, etc.). In a series of maize biparental crosses, the correlation between the observed and predicted *V*_G_ was 0.26 for yield, 0.46 for moisture, and 0.50 for test weight, with biases of only −1 to 5% in the predicted *V*_G_ (Lian et al., 2015). These correlations between predicted and observed *V*_G_ are not high enough for choosing biparental crosses on the basis of both their mean and *V*_G_.

Nearly 30 years ago, Wyman E. Nyquist and I developed a method to systematically predict *V*_G_ in biparental crosses in a line development program (Bernardo & Nyquist, 1998). This method, which was developed for a one-locus model, relied on knowing the *V*_G_ in crosses among the founder lines (referred to as founder crosses). The values of *V*_G_ in founder crosses are then used in a tabular procedure similar to that used for systematically calculating coefficients of coancestry (Emik & Terrill, 1949). Suppose A, B, and C are founder lines; D is a homozygous line derived from A/B; E is a homozygous line derived from B/C; and F is a homozygous line derived from the D(D/E) backcross population. Further suppose that the estimates of *V*_G_ in the founder crosses are 100 in A/B, 80 in A/C, and 60 in B/C (Table 1).

**Table 1.**
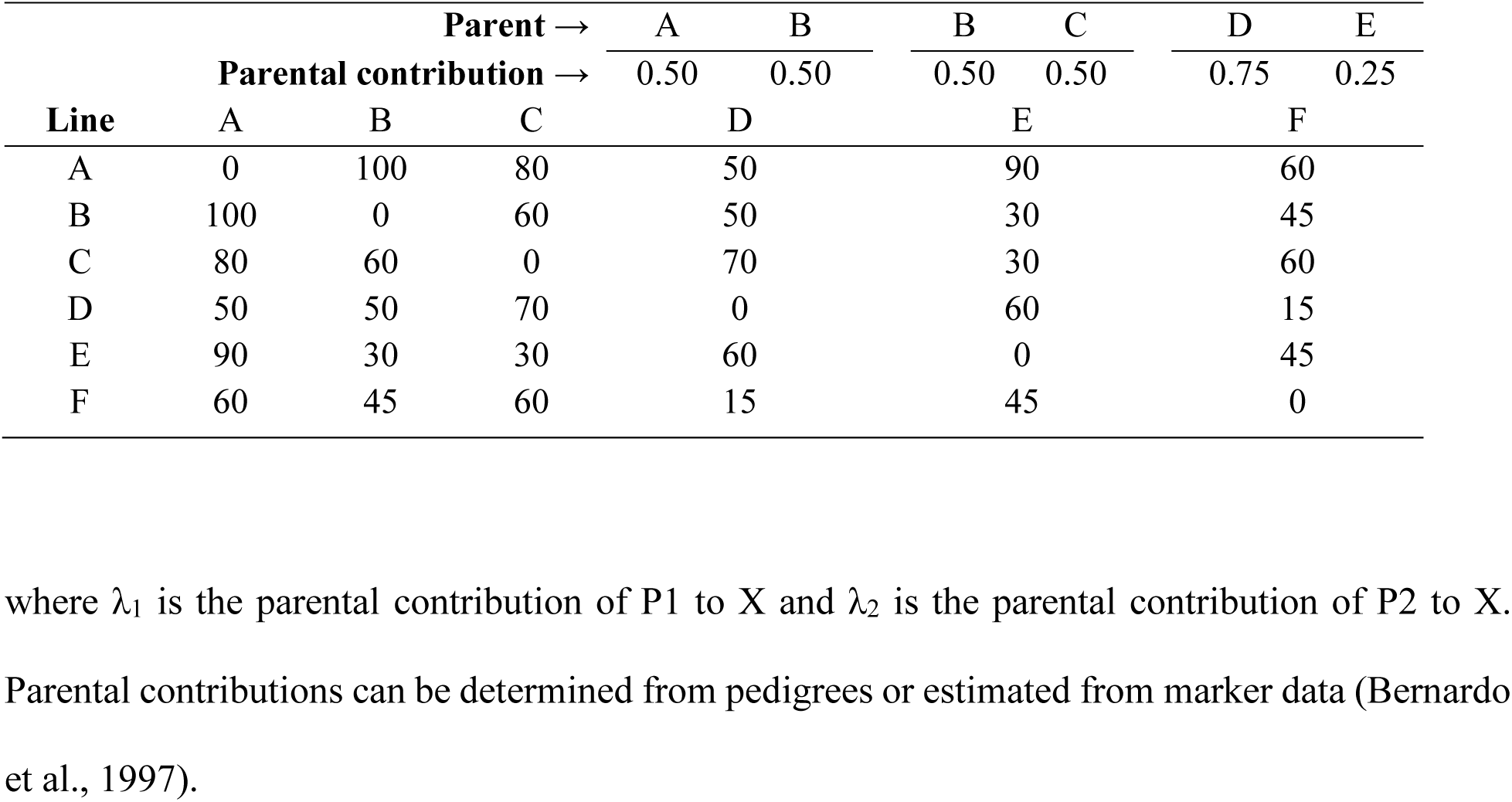
Tabular analysis of genetic variances in a hypothetical pedigree, with three founders (lines A, B, and C), two half-sib lines derived from crosses among the founders (lines D and E), and a line derived from the D(D/E) backcross (line F).

| **Parent →** |  |  |  | A | B | B | C | D | E |
|---|---|---|---|---|---|---|---|---|---|
| **Parental contribution →** |  |  |  | 0.50 | 0.50 | 0.50 | 0.50 | 0.75 | 0.25 |
| **Line** | A | B | C | D |  | E |  | F |  |
| A | 0 | 100 | 80 | 50 |  | 90 |  | 60 |  |
| B | 100 | 0 | 60 | 50 |  | 30 |  | 45 |  |
| C | 80 | 60 | 0 | 70 |  | 30 |  | 60 |  |
| D | 50 | 50 | 70 | 0 |  | 60 |  | 15 |  |
| E | 90 | 30 | 30 | 60 |  | 0 |  | 45 |  |
| F | 60 | 45 | 60 | 15 |  | 45 |  | 0 |  |

where λ<sub>1</sub> is the parental contribution of P1 to X and λ<sub>2</sub> is the parental contribution of P2 to X. Parental contributions can be determined from pedigrees or estimated from marker data (Bernardo et al., 1997).

In this example, the *V*_G_ in A/D is predicted as ½*V*_A/A_ and ½*V*_A/B_, where the subscripts refer to the parents of the biparental cross and the coefficient of ½ is the expected contribution of each parent to the genome of the progeny line. With *V*_A/A_ being zero, the predicted *V*_A/D_ is then ½*V*_A/B_ = 50 (Table 1). In general, if P1 and P2 are the parents of line X, the *V*_G_ in X/Y can be predicted as (Bernardo & Nyquist, 1998):

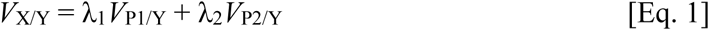

Tabular analysis involves sorting the lines from the oldest (founders A, B, and C) to the youngest (F), and systematically predicting *V*_G_ from the oldest to the most recent biparental crosses. This sorting from oldest to youngest allows the prior calculation of the *V*_G_ values needed to predict those in later biparental crosses. With F being a BC_1_-derived homozygous line, *V*_A/F_ is predicted as ¾*V*_A/D_ + ¼ *V*_A/E_, with the coefficient of ¾ being for the recurrent parent and ¼ being for the donor parent.

The Bernardo and Nyquist (1998) theoretical study had two simplifying assumptions: the quantitative trait loci (QTL) are in linkage equilibrium, and *V*_G_ is known without error in the founder crosses. When a trait is controlled by many QTL, having such QTL in a finite genome naturally leads to linkage. Sampling error is always present in estimates of *V*_G_, with the errors being large when the sizes of the founder crosses are small and heritability is low (Cockerham, 1963; Knapp et al., 1987).

Knowing the impacts of linkage and of sampling error in the estimates of *V*_G_ is therefore important if *V*_G_ in biparental crosses is to be predicted routinely via the Bernardo and Nyquist (1998) procedure. My objective in this study was to determine if *V*_G_ in biparental crosses can be systematically predicted even with linkage and with imperfect estimates of *V*_G_ in founder crosses.

## 2 MATERIALS AND METHODS

### 2.1 Genetic model

The simulated biparental crosses had 1000 polymorphic markers spaced equidistantly in a 1749 cM maize genome (Senior et al., 1996), which had 10 chromosomes. The trait was controlled by *L* = 200 QTL located at random according to a uniform distribution across the genome. Having *L* = 30 QTL was also investigated in a limited number of experiments, as described in Section 2.5.

Four founder lines, denoted by L1 to L4, were assumed. The marker loci were biallelic, as is expected for single nucleotide polymorphism (SNP) loci. The marker homozygotes were coded as 1 and –1. At each marker locus, the SNP configuration among the four founders was picked at random from six possible configurations. These configurations, which indicated the marker genotypes in L1 to L4, were as follows: (1) 1 1 –1 –1; (2) 1 –1 1 –1; (3) 1 –1 –1 1; (4) –1 1 1 – 1; (5) –1 1 –1 1; and (6) –1 –1 1 1. Each SNP locus was therefore polymorphic among the four founders and had an initial allele frequency of ½.

Each of the four founders had a different allele at each of the QTL. Similar to one of the models used by Bernardo and Nyquist (1998), the genotypic effects across the *L* QTL were normally distributed with (1) a mean of 30/√*L* in L1 1, 25/√*L* in L2, 20/√*L* in L3, and 15/√*L* in L4, and (2) a standard deviation of 1/√*L* in each of the four founder lines. The differences in the mean effects of QTL allowed *V*_G_ to vary among the founder crosses. Random *z* values were drawn from a standard normal distribution. Genotypic values were initially obtained by multiplying *z* by 1/√*L* and adding the mean that corresponded to the founder inbred.

Repulsion and coupling linkages were generated by randomly assuming one of 16 classes of loci (Dudley, 1984) among the four founder lines. These classes of loci were defined by alternating the QTL genotypes of 1 and –1 among the four founders. For example, the QTL genotypes among the four founders were 1 1 1 1 in class I; 1 1 1 –1 in class II; 1 1 –1 1 in class III; and so on until the last class (XVI), which was –1 –1 –1 –1. The final genotypic effects were calculated by multiplying the coefficients from a randomly chosen class of loci and the normally distributed genotypic effects. Suppose the randomly generated *z* value was 1.5 and class II was picked at random. In this example, the genotypic values were 1[30/√*L* + *z*(1/√*L)*] = 1[30/√200 + 1.5(1/√200)] = 2.23 in L1, 1.52 in L2, 1.17 in L3, and (– 1)[15/√200 + 1.5(1/√200)] = –1.87 in L4. Because testcross means behave in an additive manner regardless of the level of dominance (Bernardo, 2020; Hallauer & Miranda, 1981), the genotypic effects were equally valid for the performance of the lines by themselves as well as in testcrosses with the same tester.

### 2.2 Line recycling and true *V*_G_ in each biparental cross

Lines 5–10 were obtained from crosses among the founder lines. In particular, L5 was derived from L1/L2; L6 from L1/L3; L7 from L1/L4;; L8 from L2/L3; L9 from L2/L4; and L10 from L3/L4. Lines 11–25 were obtained from crosses among L5 to L10. For example, L11 was derived from L5/L6; L12 from L5/L7; and L25 from L9/L10. Finally, lines 26–40 were derived from chain crosses of L11 to L25. In particular, L26 was derived from L11/L12; L27 from L12/L13; L28 from L13/L14; and so on with the last line, L40, being derived from L25/L11.

The genotypic value of a line was calculated as the sum of genotypic values across all QTL, given the alleles carried by the line. Selection was assumed during line development in that the genotypic value of a non-founder line (L5 to L40) had to exceed the genotypic value of at least one of its parents.

Having 40 parental lines led to 40(39)/2 = 780 biparental crosses. Because of linkage disequilibrium, the *V*_G_ in each biparental cross could not be easily calculated analytically. Instead, *V*_G_ was estimated by simulating 10,000 random doubled haploids in each biparental cross and calculating the variance among the genotypic values of the doubled haploids. This estimate was then assumed equal to the true *V*_G_ in the biparental cross.

### 2.3 Estimation of *V*_G_ in founder crossses

In each of the six crosses among the four founder lines, a total of *N* = 50, 100, 200, 400, or 1000 doubled haploids were simulated. Phenotypic values of these *N* lines were then simulated in each of *N*_Env_ = 6 environments. The use of *N*_Env_ = 12 environments was also studied for a subset of experiments, as described in Section 2.5.

The phenotypic value of a line in each environment was obtained by adding an environment effect and a residual effect to the genotypic value. This residual effect was the confounded effect of genotype-environment interaction and within-environment error, assuming a single replication per environment (Bernardo, 2020; Comstock & Moll, 1963). To appropriately scale the environment and residual effects, the mean of the true *V*_G_ in the founder crosses was first calculated. The environment effects were normally distributed with a mean of zero and a variance of twice the mean true *V*_G_ in the founder crosses. The residual effects had a mean of zero and a variance scaled so that the heritability (relative to the mean true *V*_G_ in the founder crosses, but not the *V*_G_ in each cross) was *h*^2^ = 0.20, 0.50, or 0.80.

From the simulated phenotypic values of *N* lines evaluated in *N*_Env_ environments, the estimate of *V*_G_ (used in subsequent tabular analysis) in each founder cross was obtained from an analysis of variance. The observed mean squares were equated to their expectations to solve for *V*_G_.

### 2.4 Parental contributions and tabular analysis to predict *V*_G_

Markers that were polymorphic between the parents of each cross were identified. The parental contributions to progeny were estimated as the SNP similarity, for the markers polymorphic between the parents, between the first parent and the progeny line (λ_1_) and between the second parent and the progeny line (λ_2_), with λ_1_ and λ_2_ summing to 1. The 40 lines had already been sorted from oldest to youngest, and tabular analysis to predict *V*_G_ was conducted according to the procedure described by Bernardo and Nyquist (1998) and illustrated in the Introduction.

### 2.5 Statistical analysis, further experiments, and computing platform

Each simulation experiment comprised a combination of *N* (50, 100, 200, 400, and 1000) and *h*^2^ (0.20, 0.50, and 0.80). The regression of the predicted *V*_G_ on the true *V*_G_, as well as the correlation between the predicted *V*_G_ and the true *V*_G_, were calculated in each repeat. The six founder crosses, for which estimates of V*_G_* from simulated field experiments were available, were excluded from the regression and correlation analysis. Each of the 15 simulation experiments was repeated 1000 times, with the QTL locations differing in each repeat. The mean of the regression coefficient and the mean correlation were calculated across the 1000 repeats. The regression coefficients and the correlations across the 1000 repeats were sorted in ascending order, and the 25th value was considered as the 2.5th percentile value and the 975th value was considered as the 97.5th percentile value.

Five sets of additional simulations were conducted. First, the influence of the number of QTL was studied by having *L* = 30 QTL under three sets of conditions: low *N* (= 50) and *h*^2^ (= 0.20); moderate *N* (= 200) and *h*^2^ (= 0.50); and high *N* (= 1000) and *h*^2^ (= 0.80). The number of environments remained *N*_Env_ = 6. Second, the influence of the number of environments was studied by having *N*_Env_ = 12 environments, with *L* = 200 QTL and with the same three combinations of *N* and *h*^2^ mentioned above. Third, the influence of linkage was studied by assuming free recombination among loci, with *L* = 200 QTL, *N*_Env_ = 6 environments, and the same three combinations of *N* and *h*^2^ mentioned above. Fourth, the influence of sampling error in the estimates of *V*_G_ in the founder crosses was studied by conducting tabular analysis starting with the true *V*_G_ (rather than the estimated *V*_G_) in the six founder crosses. The number of QTL was *L* = 200. Fifth, the influence of selection was studied by repeating the fourth set of experiments but with the condition that the genotypic value of a non-founder line (L5 to L40) could be less than the genotypic value of both parents.

The simulations were performed via ad hoc Fortran software. The experiments ran in parallel for about 24 days on a Windows 11 computer with a 16-core, 32-thread AMD Ryzen 9 9950X processor.

## 3 RESULTS

The *V*_G_ among the six founder crosses had a mean of 274.5, and it ranged from 147.7 in the L3/L4 cross to 407.3 in the L1/L2 cross. The mean *V*_G_ among the 774 non-founder crosses was 247.6. When the true *V*_G_ was used in tabular analysis, the mean predicted *V*_G_ among the non-founder crosses was 251.6, which represented a 1.6% upward bias.

The number of lines in the founder crosses (*N*) and the mean heritability (*h*^2^) had negligible effects on the mean regression of the predicted *V*_G_ on the true *V*_G_ (denoted herein by *b*_VG_). The mean *b*_VG_ across the 1000 iterations of each simulation experiment remained almost constant at 0.51–0.52, even when *N* varied from 50 to 1000 and *h*^2^ varied from 0.20 to 0.80 (Table 2). These results were for *L* = 200 QTL and *N*_Env_ = 6 environments used in experiments to estimate *V*_G_ in the six founder crosses. As indicated by the difference between the 97.5th and 2.5th percentile values (middle 95%), the variation in *b*_VG_ across iterations was largest at lower values of *N* and *h*^2^. With *N* = 50 and *h*^2^ = 0.20, the *b*_VG_ in the middle 95% ranged from 0.02 to 1.09 (Table 2). When *N* increased to 1000, *h*^2^ had a smaller effect on the variation in *b*_VG_, with the width of the middle 95% ranging from 0.35 to 0.44.

**Table 2.** Mean regression of predicted genetic variance (*V*_G_) on true *V*_G_, and mean correlation between predicted and true *V*_G_, as influenced by the number of lines (*N*) in the founder crosses and heritability (*h*^2^) in the estimation of *V*_G_ in the founder crosses.

| ***N*** | ***h*<sup>2</sup>** | **Regression coefficient** | **Correlation** |
|---|---|---|---|
| 50 | 0.20 | 0.51 (0.02, 1.09)<sup>a</sup> | 0.49 (0.02, 0.74) |
| 100 | 0.20 | 0.52 (0.15, 0.97) | 0.56 (0.26, 0.75) |
| 200 | 0.20 | 0.52 (0.24, 0.84) | 0.63 (0.41, 0.77) |
| 400 | 0.20 | 0.51 (0.29, 0.76) | 0.66 (0.50, 0.78) |
| 1000 | 0.20 | 0.52 (0.34, 0.75) | 0.69 (0.56, 0.79) |
| 50 | 0.50 | 0.52 (0.29, 0.80) | 0.66 (0.49, 0.79) |
| 100 | 0.50 | 0.52 (0.32, 0.78) | 0.68 (0.54, 0.79) |
| 200 | 0.50 | 0.52 (0.34, 0.73) | 0.69 (0.56, 0.79) |
| 400 | 0.50 | 0.51 (0.34, 0.70) | 0.70 (0.57, 0.80) |
| 1000 | 0.50 | 0.52 (0.36, 0.72) | 0.70 (0.58, 0.79) |
| 50 | 0.80 | 0.52 (0.33, 0.75) | 0.69 (0.55, 0.80) |
| 100 | 0.80 | 0.52 (0.34, 0.74) | 0.69 (0.57, 0.80) |
| 200 | 0.80 | 0.52 (0.36, 0.72) | 0.70 (0.57, 0.80) |
| 400 | 0.80 | 0.51 (0.36, 0.70) | 0.70 (0.58, 0.80) |
| 1000 | 0.80 | 0.52 (0.36, 0.71) | 0.70 (0.58, 0.79) |

<sup>a</sup> Mean across 1000 repeats, with 2.5th and 97.5th percentiles in parentheses. Simulation experiments were for 200 quantitative trait loci and six environments used in experiments to estimate *V*<sub>G</sub> in founder crosses.

Among the non-founder crosses, the correlation between the true *V*_G_ and predicted *V*_G_, which is denoted herein by *r*_VG_, ranged from 0.49 (*N* = 50 and *h*^2^ = 0.20) to 0.70 (*N* = 400 to 1000 and *h*^2^ = 0.50 or 0.80; Table 2). The effect of *N* on *r*_VG_ was the strongest when *h*^2^ was low. As with *b*_VG_, the variability in *r*_VG_ across the 1000 iterations was largest when both *N* (= 50) and *h*^2^ (= 0.20) were low (middle 95% from 0.02 to 0.74). The middle 95% of the correlations had a narrower width when both *N* and *h*^2^ were high.

Although the results varied across iterations, near-perfect correspondence was observed between the predicted *V*_G_ averaged across all 1000 iterations and the true *V*_G_ averaged across all 1000 iterations. To illustrate, consider the results with *N* = 200 and *h*^2^ = 0.50. In a random iteration, the *b*_VG_ was 0.49 and the *r*_VG_ was 0.70 (Figure 1). In contrast, when the regression coefficient and correlation were calculated from the predicted *V*_G_ and true *V*_G_ averaged across all 1000 iterations, the *b*_VG_ was 1.00 and the *r*_VG_ was 0.997.

**Figure 1:**
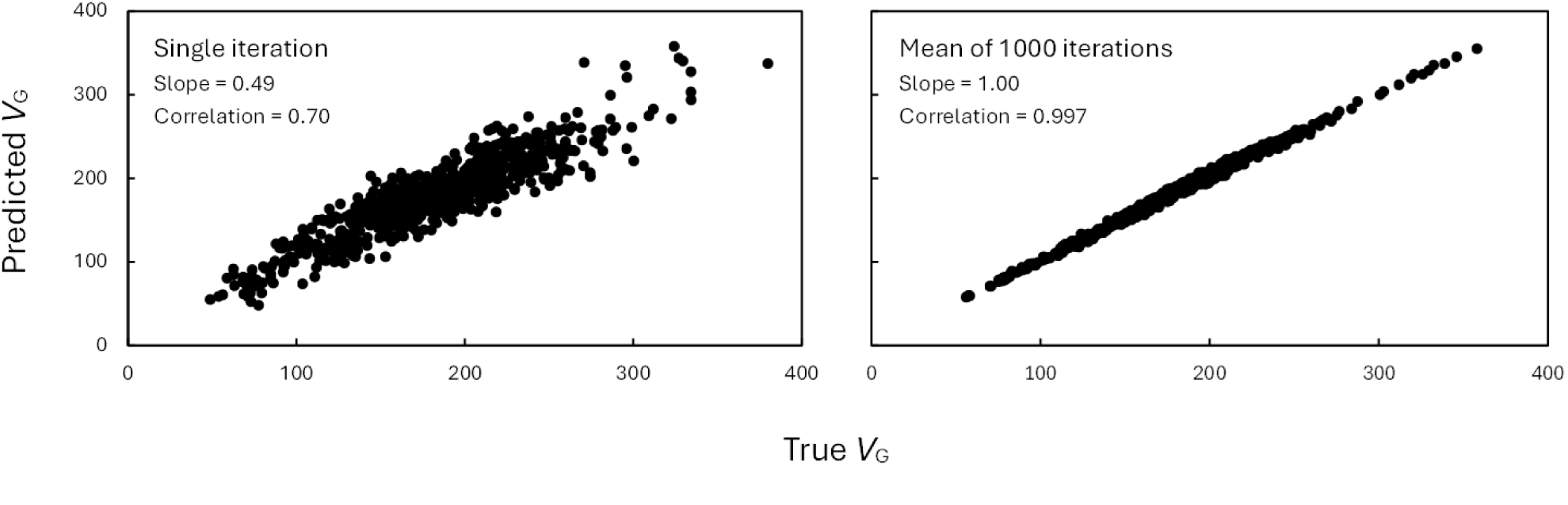
Relationship between true genetic variance (*V*_G_) and predicted *V*_G_ in a single iteration versus across 1000 iterations, with *L* = 200 quantitative trait loci (QTL), *N* = 200 lines in each founder cross, *N*_Env_ = 6 environments, and a mean heritability of *h*^2^ = 0.50. The locations of QTL across the genome differed in each iteration.

The number of environments used in estimating *V*_G_ in the six founder crosses had only a minor effect (Table 3). With *N* = 50 and *h*^2^ = 0.20, the mean *b*_VG_ increased slightly from 0.51 with *N*_Env_ = 6 (Table 2) to 0.53 with *N*_Env_ = 12 (Table 3). The number of QTL had a larger influence than did *N*_Env_. The mean *b*_VG_ decreased from 0.51–0.52 with *L* = 200 QTL (Table 2) to 0.40–0.42 with *L* = 30 QTL (Table 3). The mean *r*_VG_ decreased from 0.49–0.70 with *L* = 200 QTL (Table 2) to 0.43–0.62 with *L* = 30 QTL (Table 3).

**Table 3.**
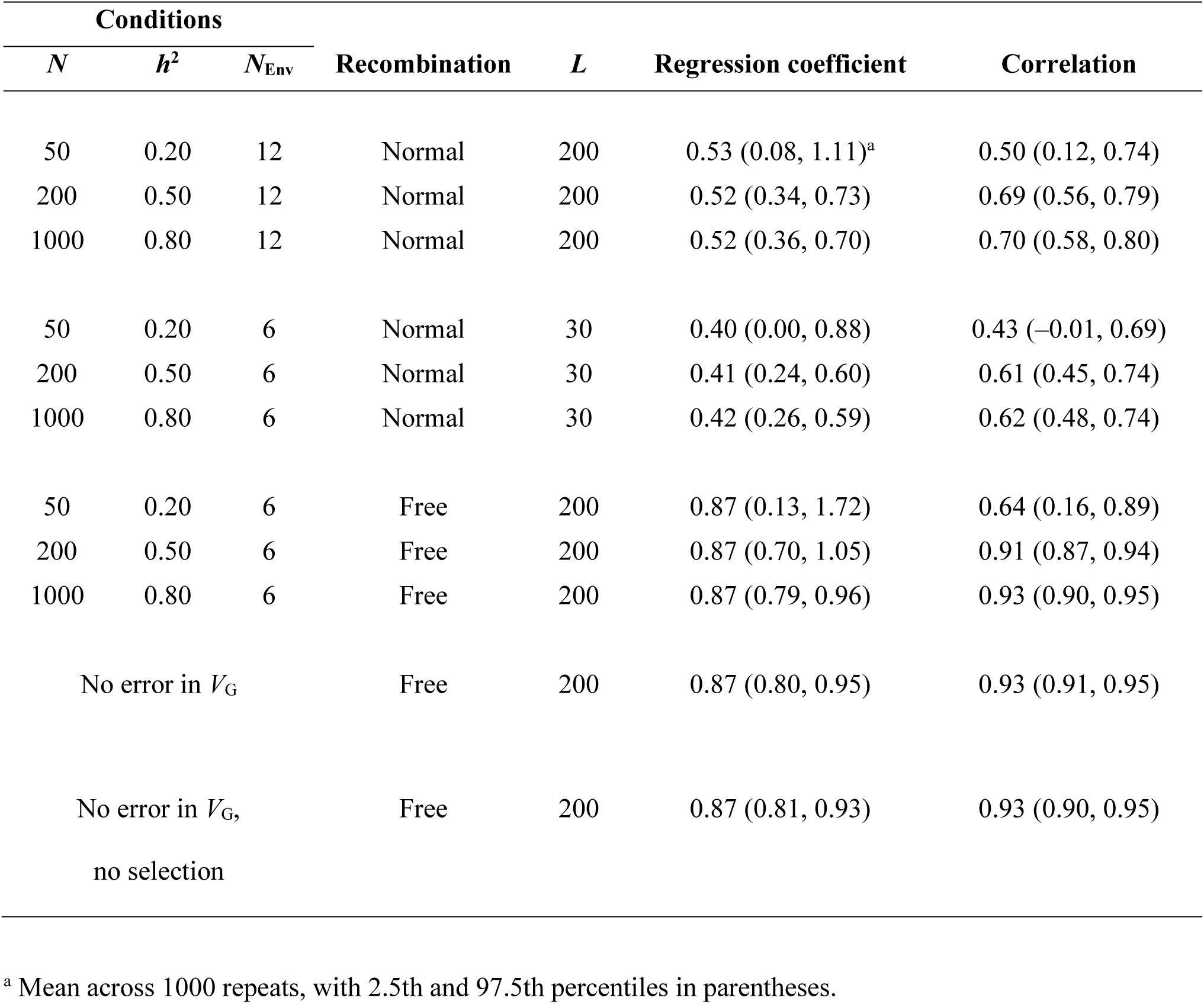
Mean regression of predicted genetic variance (*V*_G_) on true *V*_G_, and mean correlation between predicted and true *V*_G_, as influenced by the following factors: number of lines (*N*) in the founder crosses, heritability (*h*^2^), and number of environments (*N*_Env_) used to estimate *V*_G_ in the founder crosses; number of quantitative trait loci (*L*); sampling error in *V*_G_; linkage versus free recombination; and selection during line development.

| **Conditions** |  |  | **Recombination** | ***L*** | **Regression coefficient** | **Correlation** |
|---|---|---|---|---|---|---|
| ***N*** | ***h*<sup>2</sup>** | ***N*<sub>Env</sub>** |  |  |  |  |
| 50 | 0.20 | 12 | Normal | 200 | 0.53 (0.08, 1.11)<sup>a</sup> | 0.50 (0.12, 0.74) |
| 200 | 0.50 | 12 | Normal | 200 | 0.52 (0.34, 0.73) | 0.69 (0.56, 0.79) |
| 1000 | 0.80 | 12 | Normal | 200 | 0.52 (0.36, 0.70) | 0.70 (0.58, 0.80) |
| 50 | 0.20 | 6 | Normal | 30 | 0.40 (0.00, 0.88) | 0.43 (–0.01, 0.69) |
| 200 | 0.50 | 6 | Normal | 30 | 0.41 (0.24, 0.60) | 0.61 (0.45, 0.74) |
| 1000 | 0.80 | 6 | Normal | 30 | 0.42 (0.26, 0.59) | 0.62 (0.48, 0.74) |
| 50 | 0.20 | 6 | Free | 200 | 0.87 (0.13, 1.72) | 0.64 (0.16, 0.89) |
| 200 | 0.50 | 6 | Free | 200 | 0.87 (0.70, 1.05) | 0.91 (0.87, 0.94) |
| 1000 | 0.80 | 6 | Free | 200 | 0.87 (0.79, 0.96) | 0.93 (0.90, 0.95) |
| No error in *V*<sub>G</sub> |  |  | Free | 200 | 0.87 (0.80, 0.95) | 0.93 (0.91, 0.95) |
| No error in *V*<sub>G</sub>,<br>no selection |  |  | Free | 200 | 0.87 (0.81, 0.93) | 0.93 (0.90, 0.95) |

<sup>a</sup> Mean across 1000 repeats, with 2.5th and 97.5th percentiles in parentheses.

Linkage had a large effect on both *b*_VG_ and *r*_VG_. In a subset of three experiments with *L* = 200 QTL and *N*_Env_ = 6 environments, the mean *b*_VG_ increased from 0.51–0.52 with linkage (Table 2) to 0.87 with free recombination (Table 3). Likewise, the mean *r*_VG_ increased from 0.49–0.70 with linkage to 0.64–0.93 with free recombination. When free recombination was present, the use of the true *V*_G_ values instead of the most precisely estimated *V*_G_ values (*N* = 1000 and *h*^2^ = 0.80) in tabular analysis had a negligible effect, with the mean *b*_VG_ remaining 0.87 and the mean *r*_VG_ remaining 0.93. These mean values of *b*_VG_ and *r*_VG_ remained the same when selection was not imposed during line development.

## 4 DISCUSSION

### 4.1 Strengths and limitations of tabular analysis to predict *V*_G_

The results suggested that the tabular procedure developed by Bernardo and Nyquist (1998) is currently the most effective method for predicting *V*_G_. This procedure was limited by two simplifying assumptions: the QTL are unlinked, and *V*_G_ in different founder crosses is known without error. Yet even with linkage and with *V*_G_ needing to be estimated from phenotypic data, the correlations between predicted *V*_G_ and true *V*_G_ were moderately high at *r*_VG_ = 0.50–0.70 across different simulation experiments studied herein (Table 2).

When *V*_G_ was predicted from genomewide marker effects, the *r*_VG_ values obtained in previous studies at the University of Minnesota were as follows: –0.24 to 0.14 in maize (Adeyemo & Bernardo, 2019); 0.01 to 0.48 in barley (*Hordeum vulgare* L.) (Neyhart & Smith, 2019); and 0.17 to 0.88 in soybean (Wartha & Lorenz, 2024). The latter correlation of 0.88, which was for days to maturity, was driven by several plant introductions that matured about 15 days later than the elite soybean lines. The *r*_VG_ for Fusarium head blight resistance in barley was 0.61 (Tiede et al., 2015). In bread wheat (*Triticum aestivum* L.), the median correlation (across different models) between the predicted and observed genetic standard deviation ranged from 0.01 for yield to 0.59 for plant height (Oget-Ebrad et al., 2024). Caution is needed when comparing the results from a simulation study versus empirical studies. Nevertheless, the *r*_VG_ with genomewide marker effects in previous empirical studies was lower than those with tabular analysis in this simulation study.

As previously mentioned, shrinkage of genomewide marker effects toward zero has led to a downward bias when *V*_G_ was predicted from genomewide marker effects, with the downward bias approaching –100% (Neyhart and Smith, 2019). Methods to reduce this bias have been proposed (Lehermeier et al., 2017). In contrast, the mean predicted *V*_G_ with tabular analysis differed from the mean true *V*_G_ by only 1.6%.

Whereas the *r*_VG_ with tabular analysis was moderately high at 0.50–0.70, the regression of predicted *V*_G_ on true *V*_G_ was less than the desired value of *b*_VG_ = 1, with the mean observed *b*_VG_ values being about 0.50 (Table 2). As previously mentioned, the Bernardo and Nyquist (1998) method assumed linkage equilibrium and, when the simulations involved free recombination among loci, the mean *b*_VG_ increased to 0.87. This result indicated that linkage weakens the ability to predict *V*_G_ via tabular analysis.

The influence of linkage was further evidenced by the simulations with *L* = 30 QTL instead of *L* = 200 QTL. The breakage of repulsion linkages is known to release *V*_G_ that was previously hidden, whereas the breakage of coupling linkages reduces the *V*_G_ expressed in the progeny (Bernardo, 2025; Lynch & Walsh, 1998). Suppose two QTL are linked in a founder cross. If the linkage phase between the two QTL is retained in a non-founder cross, then the *V*_G_ due to the two QTL should remain unchanged. But if the linkage phase changes between a founder cross and a non-founder cross, then the *V*_G_ due to the two QTL is no longer preserved between generations. Because linkage phases are more conserved when linkage is tight, the tabular method is expected to be more effective when two QTL are tightly linked than when they are loosely linked (with free recombination being an exception). With linkage being weaker with fewer QTL, the lower *b*_VG_ with *L* = 30 QTL than with *L* = 200 QTL was therefore expected.

Even without free recombination, the results showed a near-perfect *r*_VG_ and a *b*_VG_ of 1.00 when the predicted *V*_G_ averaged across all 1000 iterations was plotted against the true *V*_G_ also averaged across all 1000 iterations (Figure 1). This result indicated that tabular analysis works extremely well when averaged over many random linkage phases. However, considering the mean *b*_VG_ and *r*_VG_ across iterations is misleading in practice because breeders will likely estimate the *V*_G_ in the same biparental population only once. As such, reducing the variability in *b*_VG_ and in *r*_VG_ is important. Results on the difference between the 97.5th percentile value and the 2.5th percentile value (middle 95%) indicated that with *h*^2^ ≥ 0.50, having *N* = 100 in each founder cross is sufficient for estimating *V*_G_ for tabular analysis. However, when *h*^2^ is lower (0.20), a few hundred progeny in each founder cross are needed to estimate *V*_G_. These numbers of progeny are consistent with those that have been recommended for estimating *V*_G_ in a plant breeding program (Bernardo, 2020; Knapp et al., 1987).

Although free recombination substantially increased the mean *b*_VG_ from about 0.50 to 0.87 (Table 3), this mean *b*_VG_ was still less than the expected value of 1. This result was attributed to inherent error in the estimates of parental contribution. As pointed out by Bernardo and Nyquist (1998), the parental contributions needed in tabular analysis are estimated from markers across the genome and are used as proxies for the parental contributions at the individual QTL themselves. Differences in the parental contributions averaged across the genome versus the parental contributions at the QTL themselves would lead to imperfect predictions of *V*_G_ via tabular analysis. With the QTL for a complex trait such as grain yield being unknown, the imperfect estimation of parental contribution at the QTL themselves will always be a limitation of tabular analysis.

Epistasis was assumed absent, and estimates of epistatic variance are typically small for different traits in different species (Bernardo, 2020; Hallauer & Miranda, 1981; Moll & Stuber, 1974). On the other hand, when two QTL exhibit epistasis, fixation at one of the two QTL can cause what was previously a small amount of epistatic variance to be converted into a larger amount of additive genetic variance (Goodnight, 1988). The tabular method relies on *V*_G_ being a linear function of parental contributions and of the *V*_G_ in prior crosses. Because epistasis distorts such linearity, it is expected to make tabular analysis of *V*_G_ less effective.

Founder crosses typically vary in *V*_G_. For example, the *V*_G_ for soybean yield was 29,700, 65,000, and 78,100 kg² ha⁻² in three founder crosses (Helms et al., 1997). Tabular analysis inherently accounts for such variation in *V*_G_ among founder crosses. Multiple alleles may be present at QTL, and the current study assumed four different alleles (one in each founder line) at each of the QTL. Having multiple alleles at QTL may contribute to variation in *V*_G_ among founder crosses and, with biallelic SNP loci, it will hinder the prediction of *V*_G_ from genomewide marker effects (Adeyemo & Bernardo, 2019) unless haplotypes are constructed to mimic multiple alleles at QTL. The use of haplotypes to model multiple alleles would be meaningful only if they capture base-level variation within each of the QTL, but this assumes that (1) SNP polymorphisms represent variation at the QTL themselves, and (2) the number of SNP loci used is large enough to capture the polymorphisms at multiple sites within a gene. These two conditions are highly unlikely in practice.

### 4.2 Application in a breeding program

Systematic prediction of *V*_G_ in a line development program will require information on *V*_G_ among biparental crosses of founder lines (Bernardo & Nyquist, 1998), as well as estimates of parental contribution to progeny from marker data (Bernardo et al., 1997). Given that marker analysis is routine for the germplasm developed in many breeding programs (Eathington et al., 2007), obtaining estimates of parental contribution should not be a limiting factor. Instead, the main limitation will be obtaining estimates of *V*_G_ in crosses among the founder lines, particularly across a range of environments sufficient for adequately sampling genotype-environment interaction effects (Comstock & Moll, 1963).

The number of possible crosses among lines quickly increases in a multiplicative fashion as the number of lines increases. Tabular analysis to predict *V*_G_ would therefore be most useful in breeding programs for which the number of founder lines (or the effective population size) is low. This criterion is likely met in commercial maize germplasm in the U.S., especially with the subdivision of germplasm into heterotic groups. The current U.S. maize germplasm that belongs to the Iowa Stiff Stalk Synthetic (BSSS) heterotic group was derived from B14, B37, B74, and to a much lesser extent B84 (Beckett et al., 2017, Nelson et al., 2008; White et al., 2020). As such, *V*_G_ would need to be estimated for only six founder crosses in the BSSS heterotic group, with *N* = 100–200 testcrosses needed to sample each of the six crosses (Table 2). Having multiple testers from the non-BSSS heterotic group complicates the logistics, and maize breeders will need to decide whether to use individual testers (and additional sets of founder testcrosses) versus a synthetic population of key non-BSSS lines as a single tester to represent the non-BSSS heterotic group. The non-BSSS heterotic group was largely derived from Mo17, Oh43, LH 82, LH123, PH207, and PHZ51 (Beckett et al., 2017, Nelson et al., 2008; White et al., 2020). The *V*_G_ would need to be estimated from 15 founder crosses in the non-BSSS heterotic group.

Specific breeding programs may also have a small number of founders, even if the germplasm base for the crop species as a whole has a larger effective population size. Much of the barley breeding germplasm at the University of Minnesota, for example, traces its lineage to only three lines: Morex, Manker, and M28 (Rasmusson & Phillips, 1997). The number of founder lines may also be small when partitioned according to maturity groups. Soybean in maturity groups 00 to IV, for example, has been estimated to have an effective population size of 11 to 15 founder lines (St. Martin, 1982). Furthermore, *V*_G_ may be predicted in subsets of non-founder crosses depending on which older, non-founder crosses have estimates of *V*_G_. In fact, *V*_G_ could be predicted on a piecemeal basis if the pair of *V*_G_ estimates needed in Eq. 1 is available.

As line development progresses from year to year, estimates of *V*_G_ will become available in non-founder crosses. These estimates of *V*_G_ in non-founder crosses may help account for the influence of linkage on *V*_G_ in subsequent crosses. Furthermore, it remains to be seen whether the availability of *V*_G_ estimates in non-founder crosses can compensate for the unavailability of *V*_G_ in some founder crosses. While tabular analysis predicts *V*_G_ from the oldest to the youngest crosses, it is also possible to predict backwards depending on which estimates of *V*_G_ have been estimated from field trials.

Consider the example given in the Introduction of P1 and P2 being the parents of line X. From Eq. 1, *V*_X/Y_ can be predicted as λ_1_*V*_P1/Y_ + λ_2_*V*_P2/Y_. However, if *V*_X/Y_ and *V*_P2/Y_ have been estimated from field trials but *V*_P1/Y_ has not, then *V*_P1/Y_ can be predicted backwards as [*V*_X/Y_ – λ_2_*V*_P2/Y_]/λ_1_. As such, it might be possible to impute the *V*_G_ in some of the founder crosses from the *V*_G_ in non-founder crosses. With estimates of *V*_G_ being available for the founder crosses as well as for a subset of the non-founder crosses, smoothing of the estimates is likely needed prior to tabular analysis. I speculate that such smoothing can be done by linear regression or by fitting linear splines (Quandt, 1958), with the latter accounting for the dissipation of linkage disequilibrium from generation to generation. Other machine learning approaches might also prove useful. These topics are worthy of future investigation.

## AUTHOR CONTRIBUTIONS

**Rex Bernardo:** Conceptualization; Data curation; Formal analysis; Investigation; Methodology; Resources; Software; Visualization; Writing-original draft; Writing-review & editing.

## Abbreviations

QTL: quantitative trait loci
SNP: single nucleotide polymorphism

